# Navigating Multi-scale Cancer Systems Biology towards Model-driven Personalized Therapeutics

**DOI:** 10.1101/2021.05.17.444410

**Authors:** Mahnoor Naseer Gondal, Safee Ullah Chaudhary

## Abstract

Rapid advancements in high-throughput omics technologies and experimental protocols have led to the generation of vast amounts of biomolecular data on cancer that now populates several online databases and resources. Cancer systems biology models built on top of this data have the potential to provide specific insights into complex multifactorial aberrations underpinning tumor initiation, development, and metastasis. Furthermore, the annotation of these single- or multi-scale models with patient data can additionally assist in designing personalized therapeutic interventions as well as aid in clinical decision-making. Here, we have systematically reviewed the emergence and evolution of (i) repositories with scale-specific and multiscale biomolecular cancer data, (ii) systems biology models developed using this data, (iii) associated simulation software for development of personalized cancer therapeutics, and (iv) translational attempts to pipeline multi-scale panomics data for data-driven *in silico* clinical oncology. The review concludes by highlighting that the absence of a generic, zero-code, panomics-based multi-scale modeling pipeline and associated software framework, impedes the development and seamless deployment of personalized *in silico* multi-scale models in clinical settings.

## Introduction

In 1971, President Richard Nixon declared his euphemistic “*war on cancer*” through the promulgation of the National Cancer Act (1–4). Five decades later, with billions spent on cancer research and development, a definitive and affordable treatment for the disease still evades humankind (5). Numerous “breakthrough” treatments have gone on to exhibit adverse side effects (6, 7) that lower patients’ quality of life (QoL) or have reported degrading efficacies(8). At the heart of this problem lies our limited understanding of the bewildering multifactorial biomolecular complexity as well as patient-centricity of cancer.

Recent advances in biomolecular cancer research have helped factored the system-level oncological manifestations into mutations across genetic, transcriptomic, proteomic, and metabolomic scales (9–12) that also act in concert (13, 14). Crosstalk between multiscale pathways comprising of these oncogenic mutations can further exacerbate the etiology of the disease (10,15–17). The combination of mutational diversity and interplay between the constituent pathways adds genetic heterogeneity and phenotypic plasticity in cancer cells (18–20). Hanahan and Weinburg (21, 22) summarized this heterogeneity and plasticity into “*Hallmarks of Cancer*” – a set of progressively acquired traits during the development of cancer.

Experimental techniques such as high throughput next-generation sequencing and mass spectrometry-based proteomics are now providing specific spatiotemporal cues on patient-specific biomolecular aberrations involved in cancer development and growth. The voluminous high-throughput patient data coupled with the remarkable complexity of the disease has also given impetus to data integrative *in silico* cancer modeling and therapeutic evaluation approaches (23). Specifically, scale-specific molecular insights into key regulators underpinning each hallmark of cancers are now helping unravel the complex dynamics of the disease (24) besides creating avenues for personalized therapeutics (25, 26). In this review, we will evaluate the emergence, evolution, and integration of multiscale cancer data towards building coherent and biologically plausible *in silico* models and their integrative analysis for employment in personalized cancer treatment in clinical settings.

### Scale-specific Biomolecular Data and its Applications in Cancer

Rapid advancements in molecular biology research, particularly in high-throughput genomics (27), transcriptomics (28), and proteomics (29) have resulted in the generation of big data on spatiotemporal measurements of scale-specific biomolecules (Figure 1A) in physiological as well as pathological contexts (30, 31). This vast and complex spatiotemporal data is expected to exceed 40 exabytes by 2025 (32), and is currently populating several online databases and repositories. These databases can be broadly categorized into seven salient database sub-types: biomolecules (33–35), pathways (36–38), networks (39–41), environment (42, 43), cell lines (44–47), histopathological images (48–50), and mutations, and drug (51–56) databases, which are discussed below.

**Figure 1.**
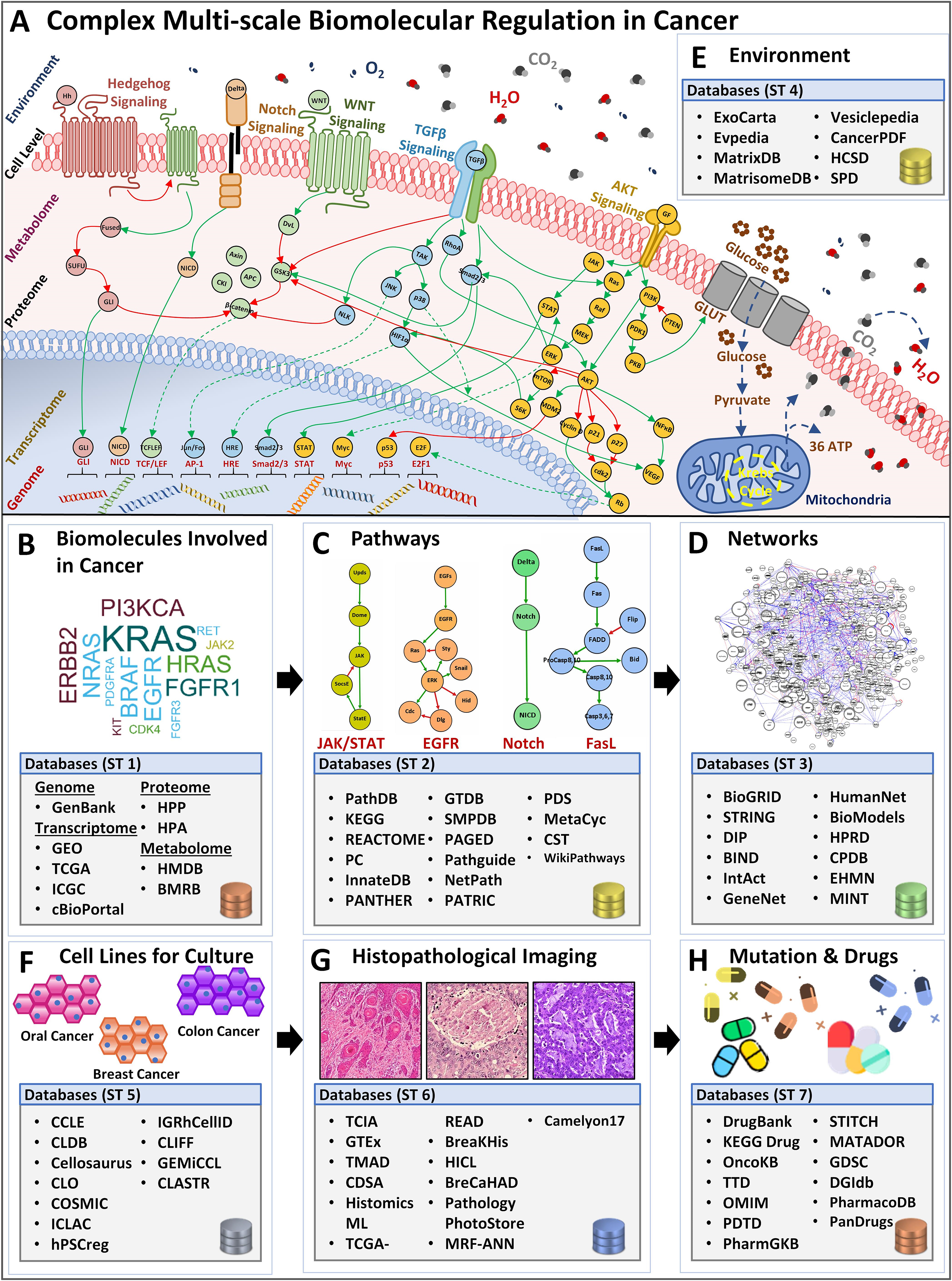
Overview of complex biomolecular regulation in cancer and scale-specific databases. **(A)** The complexity between genomic, transcriptomic, proteomic, metabolomic, cell-level, and environmental levels in a cancerous cell. Four examples of biomolecular signaling pathways are listed e.g, Hedgehog, Notch, (Wingless) Wnt, TGFβ, and AKT pathway. Stimuli from the extracellular environment signal the downstream pathway activation, in the cell, towards alternating the regulations in the proteomic, metabolomic, transcriptomic, and genomic scales, bringing out a system-level outcome in cancers. Lists **(B)** biomolecule (nucleotide, genes, proteins, and metabolites) databases such as GenBank, GEO, TCGA, HPP, HMDB, etc. **(C)** Pathways databases such as PathDB, KEGG, STRING, etc. **(D)** Networks databases such as BioGRID, DIP, BIND, etc. **(E)** Environment databases e.g, HMDB, MatrixDB, MatrisomeDB, etc. **(F).** Cell lines databases such as CCLE, CLDB, Cellosaurus, etc. **(G)** Histopathological image database, for instance, TCIA, GTEx, TMAD, etc, and **(H)** Mutation and drug databases such as DrugBank, KEGG Drug, OncoKB, etc.

#### Biomolecular Databases

Biomolecular and clinical data generated from large-scale omics approaches for cancer research can be divided into four sub-categories: (i) genome, (ii) transcriptome, (iii) proteome, and (iv) metabolome (57).

##### (i) Genome-scale Databases

The foremost endeavor to collect and organize large-scale genomics data into coherent and accessible repositories led to the establishment of GenBank in 1986(58) (Figure 1B, Supplementary Table 1). This open-access resource now forms one of the largest and most comprehensive public databases for nucleotide sequences from large-scale sequencing projects comprising over 300,000 species (59). In a salient study employing GenBank, Diez *et al.* (60) screened breast and ovarian cancer families with mutations in BRCA1 and BRCA2 genes. Medrek *et al.* (61) employed microarray profile sets from GenBank to analyze gene levels for CD163 and CD68 in different breast cancer patient groups. To date, GenBank remains the largest and most comprehensive nucleotide database; however, its data heterogeneity poses a significant challenge in its employment in the development of personalized cancer therapeutics. Towards an improved data stratification and retrieval of genome-scale data, in 2002, Hubbard *et al.* (62) launched the Ensemble genome database. Ensemble provides a comprehensive resource for human genome sequences capable of automatic annotation and organization of large-scale sequencing data. Amongst various genome-wide studies utilizing Ensemble, Easton *et al.* (63) employed the database to identify novel breast cancer susceptibility loci.

##### (ii) Transcriptome-scale Databases

Although gene-level information can facilitate the development of personalized cancer models, however, gene expression varies from cell to cell. As a result, different cancer patients have divergent genetic signatures and transcript-level gene expression(64). Therefore, high-throughput transcriptomic data has the potential to provide valuable insights into the transcriptomic complexity in cancer cells and can be useful in investigating cell state, physiology, and relevant biological events (65, 66)(Figure 1B, Supplementary Table 1). Towards developing a transcript information resource, in 2000, Edgar *et al.* (35, 67) launched the Gene Expression Omnibus (GEO) initiative. GEO acts as a tertiary resource providing coherent high throughput transcriptomic and functional genomics data. The platform now hosts over 3800 genomic datasets and is expanding exponentially and was employed by Chakraborty *et al.* (68) to annotate chemo-resistant cell line models which helped investigate chemoresistance and glycolysis in ovarian cancers. Furthermore, patient-specificity (13, 14) and mutational diversity (23) in cancer manifests across spatiotemporal scales. Hence, the availability of patient-specific data for each type of cancer has the potential to furnish valuable insights into the biomolecular foundation of the disease. In an attempt to provide cancer type-specific mutation data, Wellcome Trust’s Sanger Institute (69) developed Catalogue of Somatic Mutations in Cancer (COSMIC) (69) comprising of 10,000 somatic mutations from 66,634 clinical samples. Through the employment of COSMIC, Schubbert *et al.* (70) decoded Ras activity in cancers and developmental disorders, while Weir *et al.* (71) characterized the genome of lung adenocarcinomas. The curation of patient-specific gene and protein expression data then led to the development of The Cancer Genome Atlas (TCGA) (72). TCGA also captures the copy number variations and DNA methylation profiles for different cancer subtypes. TCGA’s potential (53, 73) is well exhibited by Leiserson *et al.*’s (74) investigation that helped identify combinations of rare somatic mutations across pathways and protein complexes. Davis *et al.* (75) further evaluated the genomic landscape of chromophobe renal cell carcinomas (ChRCCs) to elicit molecular patterns as clues for determining the origin of cancer cells. To facilitate data management across different cancer projects as well as to ensure data uniformity towards developing data-driven models, the International Cancer Genome Consortium (ICGC) was launched in 2010 (76). ICGC adopts a federated data storage architecture that enables it to host a collection of scale-specific data from TCGA and 24 other projects (77). Burn *et al.* (78) estimated the distribution of cytosine in liver tumor data using ICGC, while Supek *et al.* (79) compared mutation rates between different human cancers. Although numerous databases have been established to store large-scale genomic data, however, insights from an integrated analysis of genomic data across databases have the potential to provide precise biomolecular cues into complex processes and evolution in cancer. Such capability was provided by cBio Cancer Genomics Portal (cBioPortal) (80) in 2012, with multidimensional dataset retrieval, and exploration from multiple databases. The platform additionally provides data visualization tools, pathway exploration, statistical analysis, and selective data download features for seamless utilization of large-scale genomics data across genes, patient samples, projects, and databases (81). Numerous studies have effectively employed cBioPortal (82–84), in particular, Jiao *et al.* (85) evaluated the prognostic value of TP53 and its correlation with EGFR mutations in advanced non-small-cell lung cancer (NSCLC). Hou *et al.* (86) also used cBioPortal to deduce targetable genotypes which are present in young patients with lung adenocarcinomas.

##### (iii) Proteome-scale Databases

Gene transcriptomics is limited in providing a deterministic proteome profile (87–89). Particularly, transcripts produced in a cell can be degraded, translated inefficiently, or modified due to post-translational modification (90, 91) resulting in no or a very small amount of functional protein (66). This relatively low correlation between transcriptome and proteome data was highlighted in 2019, by Bathke *et al.* (92) where it was shown that an increase in transcripts synthesis cannot be directly associated with an increase in functional response in a cell. To facilitate functional analysis, there is a need to utilize proteomic-level data, which can help to capture a more accurate quantitative assessment of complex biomolecular regulations for functional studies (Figure 1B, Supplementary Table 1). Following the successful completion of the Human Genome Project (HGP) (1998), in 2003, a group of Swedish researchers reported the Human Protein Atlas (HPA) (93, 94) with an aim to map the entire set of human proteins in cells, tissues, and organs for normal as well as cancerous state (95). HPA employs large-scale omics-based technologies to localize and quantify protein expression patterns. The database has successfully managed to host comprehensive information on human proteins from cells, tissues, pathology, brain, and blood regions-related studies. HPA data can be employed for various purposes such as investigating the spatial distribution of proteins in different tissue and comparing normal and cancerous protein expression patterns across samples, etc (93). In a salient study employing HPA, Gámez-Pozo *et al.* (96) localized the expression pattern of proteins to aid in the profiling of human lung cancer subtypes, while Imberg-Kazdan *et al.* (97) identified novel regulators of androgen receptor function present in prostate cancer. Another significant stride towards proteome-level information came with the establishment of the Human Proteome Project (HPP) (98, 99) in 2008 (100) by the Human Proteome Organization (HUPO). HPP consolidated mass spectrometry-based proteomics data, and bioinformatics pipelines, with the aim to organize and map the entire human proteome. To date, numerous studies have utilized HPP towards identifying the complex protein machinery involved in cancer cell fate outcomes (101–106). Amongst the foremost attempts, in 2001, Sebastian *et al.* (107) employed the HPP platform to deduce the complex regulatory region of the human CYP19 gene (‘*armatose’*), one of the contributors to breast cancer regulation. HPP project was later segmented into “biology and disease-oriented HPP” (B/D HPP) (108) and chromosome-centric HPP (C-HPP) (109). Gupta *et al*. (110) carried out an extensive analysis of existing experimental and bioinformatics databases to annotate and decipher proteins associated with glioma on chromosome 12, while, Wang *et al.* (111) performed a qualitative and quantitative assessment of human chromosome 20 genes in cancer tissue and cells using C-HPP resources.

##### (iv) Metabolome-scale Databases

Metabolic reprogramming is one of the earliest manifestations during tumorigenesis (112) and, therefore, potentiates identification of biomarkers involved in cancer onset, its prognosis as well as treatment. Large-scale efforts to collect metabolomics data led to the development of several online databases (113–115) (Figure 1B, Supplementary Table 1) including the Golm Metabolome Database (GMD) (113), in 2004. GMD provides a comprehensive resource on metabolic profiles, customized mass spectral libraries, along with spectral information for use in metabolite identification. GMD was employed in 2011 by Wedge *et al*. (116) to identify and compare metabolic profiles in serum and plasma samples for small-cell lung cancer patients towards determining optimal agent for onwards analysis. Whereas, in 2013, Pasikanti *et al*. (117) utilized GMD to identify biomarker metabolites present in bladder cancer. To allow for large-scale metabolic data stratification and retrieval, in 2007, Wishart et al. published the Human Metabolome Database (HMDB) (114). HMDB contains organism-specific information on metabolites across various biospecimens and their accompanying environments. It is now the world’s largest metabolomics database with around 114,100 metabolites that have been characterized and annotated. HMDB was employed by Sugimoto *et al.* (118) to identify environmental compounds specific to oral, breast, and pancreatic cancer profiles, while Agren *et al.* (119) constructed metabolic network models for 69 human cells and 16 cancer types. Although HMDB supports data deposition and dissemination, however, integrated exploratory analysis is not available. The Metabolomics Workbench (115), reported in 2016, provides information on metabolomics metadata and experimental data across species, along with an integrated set of exploratory analysis tools. The platform also acts as a resource to integrate, deposit, track, analyze, as well as disseminate large-scale heterogeneous metabolomics data from a variety of studies. In a case study built using this platform, Hattori *et al*. (120) studied cancer progression by metabolic reprogramming in myeloid leukaemia, whereas, Playdon (121) investigated dietary biomarkers in breast cancer etiology.

#### Biomolecular Pathway Databases

Investigations restricted to single specific biomolecular scales have limited translational potential as cancer dysregulation is driven by tightly coupled biomolecular pathways constituted by biomolecules from a variety of spatiotemporal scales (discussed above). Such biomolecular pathways represent organized cascades of interactions integrating different spatiotemporal scales towards reaching specific phenotypic cell fate outcomes. Numerous scale-specific and multi-omics biomolecular pathway databases now exist to help retrieve, store and analyze existing pathway information towards understanding cellular communication in light of complex cancer regulation (38,122,123). One of the earliest attempts at integrating genomics data for pathway construction was in 1995, with the establishment of the Kyoto Encyclopedia of Genes and Genomes (KEGG) database (122) (Figure 1C, Supplementary Table 2). Over time, KEGG has significantly expanded to include high-throughput multi-omics data (124). As a result, the resource is divided into fifteen sub-groups including KEGG Genome (for genome-level pathways), KEGG Compound (for small molecules level pathways), KEGG Gene (for gene and protein pathways), KEGG Reaction (for biochemical reaction and metabolic pathways), etc. Li *et al.* (125) used the KEGG database to perform pathway enrichment analysis for predicting the function of circular RNA (circRNA) dysregulation in cancer, while Feng *et al.* (126) identified genes associated with poor prognosis in ovarian cancer. To furnish information on pathways limited to gene-protein and protein-protein interactions for high-throughput functional analysis studies, PANTHER (protein annotation through evolutionary relationship) database was established in 2010 (38, 123). PANTHER hosts information on ontological gene and protein-protein interaction pathways by leveraging GenBank and Human Gene Mutation Database (HGMD) (127), etc. Turcan *et al.* (128) employed PANTHER to perform network pathway enrichment for biological processes in differentially expressed genes to investigate IDHI mutations in glioma hypermethylation phenotype. In order to store metabolic pathway information in a cell, Karp *et al.* (129) developed MetaCyc, a comprehensive reference database of metabolic pathways. MetaCyc is currently available as a web-based resource with metabolic pathway information which can be employed to investigate metabolic reengineering in cancers, carry out biochemistry-based studies, and explore cancer cell metabolism, etc. Tang and Aittokalli (130) demonstrated the utility of MetaCyc database by annotating a human metabolic network to evaluate multi-target cancer treatments.

#### Biomolecular Network Databases

The regulatory complexity of cancer is compounded by the cross-talk between multi-scale biomolecular pathways resulting in the formation of interaction networks (Figure 1D, Supplementary Table 3). One of the earliest biomolecular network databases, the Biomolecular Interaction Network Database (BIND) (131) was established in 2001, with an aim to organize biomolecular interactions between genes, transcripts, proteins, metabolites, as well as small molecules. Chen *et al.* (132) employed BIND to construct a biological interaction network (BIN) towards investigating tyrosine kinase regulation in breast cancer development. Although BIND provides a comprehensive resource of predefined interacting pathways, however, it does not contain ‘indirect’ interaction information. In contrast, the Molecular INTeraction database (MINT) (41), developed in 2002, curates existing literature to develop networks with both direct as well as indirect interactions from large-scale projects with information from genes, transcripts, proteins, promoter regions, etc. MINT can store data on “functional” interaction as well such as enzymatic properties and modifications present in a biomolecular regulatory network. The database was employed by Sun *et al.* (133) to construct a human interactome network to evaluate cancer patients’ interactome, whereas, Vinayagam *et al.* (134) constructed an HIV (human immunodeficiency virus) network that helped identify novel cancer genes across genomic datasets. The Database of Interacting Proteins (DIP) (135) was developed to mine existing literature and experimental studies on biomolecules and their pathways to construct interacting protein networks. Numerous studies on cancer research have been reported using DIP (136–138). Goh *et al.* (136) constructed a protein-protein interaction network for investigating liver cancer using network information from DIP while Zhao *et al.* (137) identified microRNA-regulated autophagy pathways in plants. To further consolidate and integrate protein interaction data across pathways as well as organisms, the Search Tool for the Retrieval of Interacting Genes/Proteins – STRING database was developed in 2005 (139). STRING provides a comprehensive text-mining and computational prediction platform which is accessible through an intuitive web interface (140). STRING database also provides interaction weights for edges between biomolecules to show an estimated likelihood for each interaction in the network (140). Mlecnik *et al.* (141) employed the STRING database to study T-cells homing factors in colorectal cancer. Later, Bindea *et al.* (142) assembled a gene-gene correlation network to investigate natural immunity to cancer in humans.

#### Environment Databases

Each pathway within a biomolecular network requires input cues from the extracellular environment for onward downstream signal transduction (143–145). In the case of cancer, the biomolecular milieu constituting the tumor microenvironment (TME) acts as a niche for tumor development, metastasis, as well as therapy response (146) (Figure 1E, Supplementary Table 4). Efforts to curate information from the environmental factors such as metabolites, matrisome, and other environmental compounds led to the development of MatrixDB (42) in 2011, which curated matrix-based information on interactions between extracellular proteins and polysaccharides. MatrixDB additionally links databases with information on genes encoding extracellular proteins such as Human Protein Atlas (147) and UniGene (148) as well as host transcripts information. Celik *et al.* (149) employed MatrixDB data to evaluate Epithelial-mesenchymal transition (EMT) inducers in the environment in nine ovarian cancer datasets. With an aim to host studies on extracellular matrix (ECM) proteins from normal as well as disease-inflicted tissue samples, MatrisomeDB (43) was established in 2020. The database contains 17 studies on 15 physiologically healthy murine and human tissue, and 6 cancer types from different stages (including breast cancers, colon cancers, ovarian cancers, lung cancers, and melanomas) along with other diseases. Levi-Galibov *et al.* (150) employed the MatrisomeDB to investigate the progression of chronic intestinal inflammation to colon cancer.

#### Cell Line Databases

*In vitro* cell lines derived from cancer patients have become an essential tool for clinical and translational research (151). These cell lines are defined based on gene expression profiles and morphological features which have been cataloged in various databases such as the Cancer Cell Line Encyclopedia (CCLE) (46) (Figure 1F, Supplementary Table 5). CCLE contains mutation data of 947 different human cancer cell lines coupled with pharmacological profiles of 24 anti-cancer drugs (46) for evaluating therapeutic effectiveness and sensitivity. Li *et al.* (152) employed CCLE data to investigate cancer cell line metabolism, while Hanniford *et al.* (153) demonstrated epigenetic silencing of RNA during invasion and metastasis in melanoma. Other cell line databases include Cell Line Data Base (CLDB) (47), and The COSMIC Cell Lines Project (44), and CellMinerCDB (45). The CellMinerCDB (2018) curates data from National Cancer Institue (NCI) (154), BROAD institute (155), Sanger institute (156), and Massachusetts General Hospital (MGH) (157) and provides a platform for pharmacological and genomic analysis.

#### Histopathological Image Databases

Additionally, histopathological image datasets derived from the microscopic examination of tumor biopsy samples furnish information on cellular structure, function, chemistry, morphology, etc. Numerous histopathological image-based databases have been developed to store, manage and retrieve such information (Figure 1G, Supplementary Table 6). Amongst these databases, The Cancer Imaging Archive (TCIA) (48), reported in 2013, provides a multi-component architecture with various types of images including region-specific (e.g., Breast), cancer-type specific (e.g., TCGA-GBM, TCGA-BRCA), radiology, and anatomy images (e.g., Prostate-MRI). The cancer image collection in TCIA is captured using a variety of modalities including radiation treatment, X-ray, mammography, and computed tomography (CT), etc (158). Zhu *et al.* (159) exhibited the value of TCIA by predicting the risk for breast cancer recurrence, while Sun *et al.* (160) employed image data to perform a cohort study to validate a radiomics-based biomarker in cancer patients. In 2013, Image data from TCIA was integrated with The Cancer Digital Slide Archive (CDSA) (49) which hosts imaging and histopathological data and provides more than 20,00 images from 22 different cancers. The integrated datasets in CDSA demonstrate the clinical association of genomic data with pathology imaging. The whole-slide images of individual patients are also reported in CDSA, thereby linking tumor morphology with the patient’s genomic and clinical data. Khosravi *et al.* (161) performed a deep convolution study, using CDSA, to distinguish heterogeneous digital pathology images across different types of cancers. To associate a relationship between a patient’s genetic information and histology images Genotype-Tissue Expression (GTEx) (50) was reported in 2014. The GTEx project was created from datasets from 1000 patients containing tissue-specific information on gene expression and regulation.

#### Mutation and Drug Databases

Pharmacological investigations have elucidated the mechanism and efficacies of numerous cancer drugs, in clinical and preclinical studies (162, 163). Databases with drug-target information can be employed in precision oncology towards designing efficacious patient-centric therapeutic strategies (Figure 1H, Supplementary Table 7). Such databases include DrugBank (164), established in 2006, which contains information from over 4100 drug entries, 800 FDA-approved small molecules, and 14,000 protein or drug target sequences. DrugBank combines drug data with drug-target information this, in turn, can provide potential cues for various applications in cancer biology including *in silico* drug target discovery, drug design, drug interaction prediction, etc. In a study employing DrugBank, Augustyn *et al.* (165) evaluated potential therapeutic targets of ASCL2 genes in lung cancers, while Han *et al.* (166) determined synergistic combinations of drug targets in K562 chronic myeloid leukemia (CML) cells to dissect functional genetic interaction (GI) networks. Further to evaluate drugs in light of the patient’s genomic signature, the PanDrugs (56) database was established in 2019. The platform currently hosts data from 24 sources and 56297 drug-target information along with 9092 unique compounds and 4804 genes. Fernández-Navarro *et al.* (167) prioritized personalized drug treatments using PanDrugs, for Acute T-cell lymphoblastic leukemia (T-ALL) patients.

Altogether, the availability of voluminous high-resolution biomolecular data has enabled the development of a quantitative understanding of aberrant mechanisms underpinning hallmarks of cancer as well as create avenues for personalized therapeutic insights.

### Data-driven Integrative Modeling in Cancer Systems Biology

The need to prognosticate system-level outcomes in light of oncogenic dysregulation (168, 169) has led researchers to develop integrative data-driven computational models (170–177). Such models can help decode emergent mechanisms underpinning tumorigenesis as well as aid in the development of patient-centered therapeutic strategies (170–177). These *in silico* models can be broadly grouped into four salient sub-scales as biomolecular (178–180), environment (181–183), cell level (184, 185), and multi-scale cancer models (186–188) (Figure 2).

**Figure 2.**
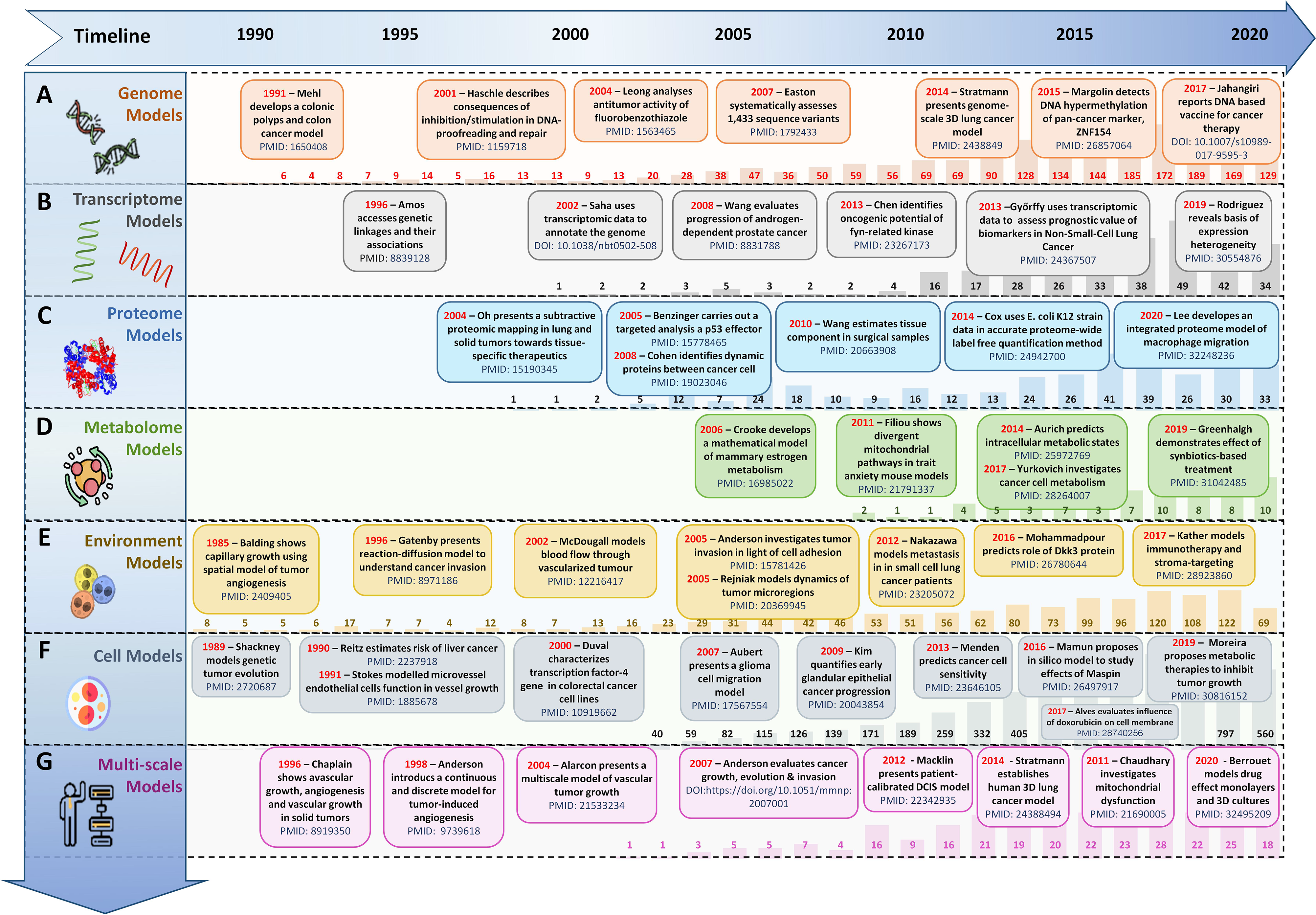
Evolution timeline of *in silico* scale-specific and multi-scale data-drive cancer models. Timeline of salient *in silico* scale-specific and multi-scale cancer models, along with PubMed yearly report (1990–2020) to display the evolutionary trends seen in the development of **(A)** genome-scale cancer models, **(B)** Transcript-level cancer models, **(C)** Proteome-scale models, **(D)** Metabolome scale models, **(E)** Environment-based models, **(F)** Cell-level models, and **(G)** Multi-scale cancer models.

#### Biomolecular-scale Models

*In silico* biomolecular models of cancer can be classified into (i) genome-scale, (ii) transcriptome-scale, (iii) proteome-scale, (iv) metabolome-scale models.

##### (i) Genome-scale Models

Amongst initial attempts at developing cancer gene regulation models, in 1987, Leppert *et al.* (178) developed a computational model to study the genetic locus of familial polyposis coli and its involvement in colonic polyposis and colorectal cancer. The study was also validated by Mehl *et al.* (179) in 1991, which further elucidated the formation and development of familial polyposis coli genes in colorectal cancer patients. In 2014, Stratmann *et al.* (180) developed a personalized genome-scale 3D lung cancer model to study epithelial-mesenchymal transition (EMT) by TGFβ-based stimulation, while in 2016, Margolin *et al.* (189) developed a blood-based diagnostic model to help detect DNA hypermethylation of potential pan-cancer marker ZNF154. In 2017, Jahangiri *et al.* (190) employed an *in silico* pipeline to evaluate Staphylococcal Enterotoxin B for DNA-based vaccine for cancer therapy. A similar genome-scale model of DNA damage and repair was proposed by Smith *et al.* (191) to evaluate proton treatment in cancer (Figure 2A).

##### (ii) Transcriptome-scale Models

*In silico* models developed using transcriptomic expression data can assist in comparing gene expressions in cancer for investigating genetic heterogeneity in tumorigenesis and cancer development (192) as well as towards precision therapy (193) (Figure 2B). In 2003, Huang *et al.* (194) developed a mathematical model using a large-scale transcriptional dataset of breast cancer patients to elucidate patterns of metagenes for nodal metastases and relapses. In another large-scale cancer transcriptomics study, 3,000 patient samples from nine different cancer types were used to decode the genomic evolution of cancer by Cheng *et al.* (195). In 2013, Chen *et al.* (196) employed an *in silico* pipeline which helped identify 183 new tumor-associated gene candidates with the potential to be involved in the development of hepatocellular carcinoma (HCC), while, in 2014, Agren *et al.* (197) reported a personalized transcriptomic data-based model to identify anticancer drugs for HCC. In 2019, Béal *et al.* (198) reported a logical network modeling pipeline for personalized cancer medicine using individual breast cancer patients’ data using existing gene expression databases. The pipeline was validated in 2021 (199), using *in silico* personalized logical models to investigate melanomas and colorectal cancers in response to BRAF treatments. In a similar study conducted in 2019, Rodriguez *et al.* (200) developed a mathematical model for breast cancer using transcriptional regulation data to predict hypervariability in a large dynamic dataset and reveal the basis of expression heterogeneity in breast cancer.

##### (iii) Proteome-scale Models

To capture the quantitative aspects of biomolecular regulations and for functional studies (88, 89), data-driven proteomic-based cancer models are essential (Figure 2C). Such models can be particularly helpful in diagnostic as well as prognostic purposes as well as for monitoring response to treatment (88, 89). In a study employing proteome-level information, in 2011, Baloria *et al.* (201) carried out an *in silico* proteome-based characterization of the human epidermal growth factor receptor 2 (HER-2) to evaluate its immunogenicity in an *in silico* DNA vaccine. Akhoon *et al.* (202) simplified this approach with the development of a new prophylactic *in silico* DNA vaccine using IL-12 as an adjuvant. In 2017, Fang *et al.* (203) further employed proteome level data towards predicting *in silico* drug-target interactions for targeted cancer therapeutics, while in 2018, Azevedo *et al.* (204) designed novel glycobiomarkers in bladder cancer. Recently, in 2020, Lee *et al.* (205) have reported an integrated proteome model of macrophage migration in a complex tumor microenvironment. However, proteome-level models are limited in their ability to provide a complete analysis of the biomolecules present in a cell since they lack information on low molecular weight biomolecular compounds such as metabolites (112).

##### (iv) Metabolome-scale Models

Metabolic data-driven cancer models can be especially useful in understanding cancer cell metabolism, mitochondrial dysfunction, metabolic pathway alteration, etc (Figure 2D). In 2007, Ma *et al.* (206) developed the Edinburgh Human Metabolic Network (EHMN) model with more than 3000 metabolic reactions alongside 2000 metabolic genes for employment in metabolite-related studies and functional analysis. In 2011, Folger *et al.*’s (207) employed the EHMN model to propose a large-scale flux balance analysis (FBA) model for investigating metabolic alterations in different cancer types and for predicting potential drug targets. In 2014, Aurich *et al.* (208) reported a workflow to characterize cellular metabolic traits using extracellular metabolic data from lymphoblastic leukemia cell lines (Molt-4) towards investigating cancer cell metabolism. Yurkovich *et al.* (209) augmented this workflow in 2017 and reported eight biomarkers for accurately predicting quantitative metabolite concentration in human red blood cells. Alakwaa *et al.* (210) employed a mathematical model to predict the status of Estrogen Receptor in breast cancer metabolomics dataset, while in 2018, Azadi *et al.* (211) used an integrative *in silico* pipeline to evaluate the anti-cancerous effects of *Syzygium aromaticum* employing data from the Human Metabolome Database (114).

#### Environment Models

Integrative mathematical models of environmental cues and extracellular matrix can help researchers abstract tumor microenvironments. Such data-driven models can be used to study angiogenesis (212), cell adhesion (213), and vasculature (214) (Figure 2E). Amongst initial attempts at developing cancer environment-based *in silico* models, in 1972, Greenpan *et al.* (181) developed a solid carcinoma *in silico* model to evaluate cancer cell behavior in limited diffusion settings. In 1976, the model was expanded (182) to investigate tumor growth in asymmetric conditions. In 1996, Chaplain developed a mathematical model to elucidate avascular growth, angiogenesis, and vascular growth in solid tumors (183). Anderson and Chaplain (215), in 1998, expanded this strategy and reported a continuous and discrete model for tumor-induced angiogenesis. This modeling approach was further augmented in 2005 by Anderson (216) to a hybrid mathematical model of a solid tumor to study cellular adhesion in tumor cell invasion. Organ-specific metastases and associated survival ratios in small cell lung cancer patients have been modeled and evaluated (217) using similar models.

#### Cell Models

To model cell population-level behavior in cancer, researchers are increasingly developing innovative cell line *in silico* models which can complement *in vivo* wet-lab experiments, while overcoming wet-lab limitations (218) (Figure 2F). Such models are employed to investigate cell-to-cell interactions and evaluate physical features of the synthetic extracellular matrix (ECM) (218), etc (184, 219). Amongst initial attempts at developing *in silico* cell line models, in 1989, Shackney *et al.* (184) reported an *in silico* cancer cell model to study tumor evolution. Results showed an association between discrete aneuploidy peaks with the activation of growth-promoting genes. In 2007, Aubert *et al.* (219) developed an *in silico* glioma cell migration model and validated cell migration preferences for homotype and heterotypic gap junctions with experimental results. Gerlee and Nelander (185) expanded this work, in 2012, to investigate the effect of phenotype switching on glioblastoma growth. Although cell-based cancer models helped provide scale-specific insights into cancer, however, they remain limited in investigating the spatiotemporal tissue diversity and heterogeneity in cancer patients.

#### Multi-scale Models

Recently, data-driven multi-scale models are becoming increasingly popular in cancer (220) (Figure 2G). One of the earliest attempts at developing multi-scale cancer models was in 1985 when Balding developed a mathematical model to demonstrate tumor-induced capillary growth (212). In 2000, Swanson *et al.* (221) proposed a quantitative model to investigate glioma cells. In a similar study, Zhang *et al.* (222) generated a 3D, multiscale agent-based model of the brain to study the role (EGFR)-mediated activation of signaling protein phospholipase role in a cell’s decision to either proliferate or migrate. In 2010, Wang *et al.* (187) also took a multi-scale agent-based modeling approach to identify therapeutic targets in concurrent EGFR-TGFβ signaling pathway in non-small cell lung cancer (NSCLC). Later in 2011, (188) they employed the approach to also identify critical molecular components in NSCLC. Similarly, Perfahl *et al.* (223) developed a multi-scale vascular tumor growth model to investigate spatiotemporal regulations in cancer and response to therapy. In 2007, Anderson *et al.* (224) proposed a mathematical model for studying cancer growth, evolution, and invasion. This model was later built upon by Chaudhary *et al.* (173) with a multi-scale modeling strategy to investigate tumorigenesis induced by mitochondrial incapacitation in cell death, in 2011. In 2017, Vavourakis *et al.* (225) developed a multi-scale model to investigate tumor angiogenesis and growth. In 2017, Norton *et al.* (226) used an agent-based computational model of three negative breast cancer to study the effects of migration of CCR5+ cancer cells, stem cell proliferation, and hypoxia on the system. They later (227) reported an agent-based and hybrid model to investigate tumor immune microenvironment, in 2019. In the same year, Karolak *et al.*(228) modeled *in silico* breast cancer organoid morphologies (229) to help elucidate efficacies amongst drug treatment based on the morphophenotypic classification. Similarly, Berrouet *et al.* (230) employed a multi-scale mathematical model to evaluate the effect of drug concentration on monolayers and spheroid cultures.

Summarily, data-integrative computational models have now assumed the forefront in decoding the complex biomolecular regulations involved in cancer and are increasingly been employed for the development of personalized preclinical models as well as therapeutics design (231).

### Software Platforms for Modeling in Cancer Systems Biology

Over the past decade, *in silico* modeling of cancer has gained significant popularity in systems biology research (170–177). In particular, data-drive computational models are now acting as an enabling technology for precision medicine and personalized treatment of cancer. To date, several single and multi-scale software platforms have been reported to model biomolecular (180,232–239), environmental (240), cell-level (241–243), and multi-scale (244–246) information into coherent *in silico* cancer models (Figure 3).

**Figure 3.**
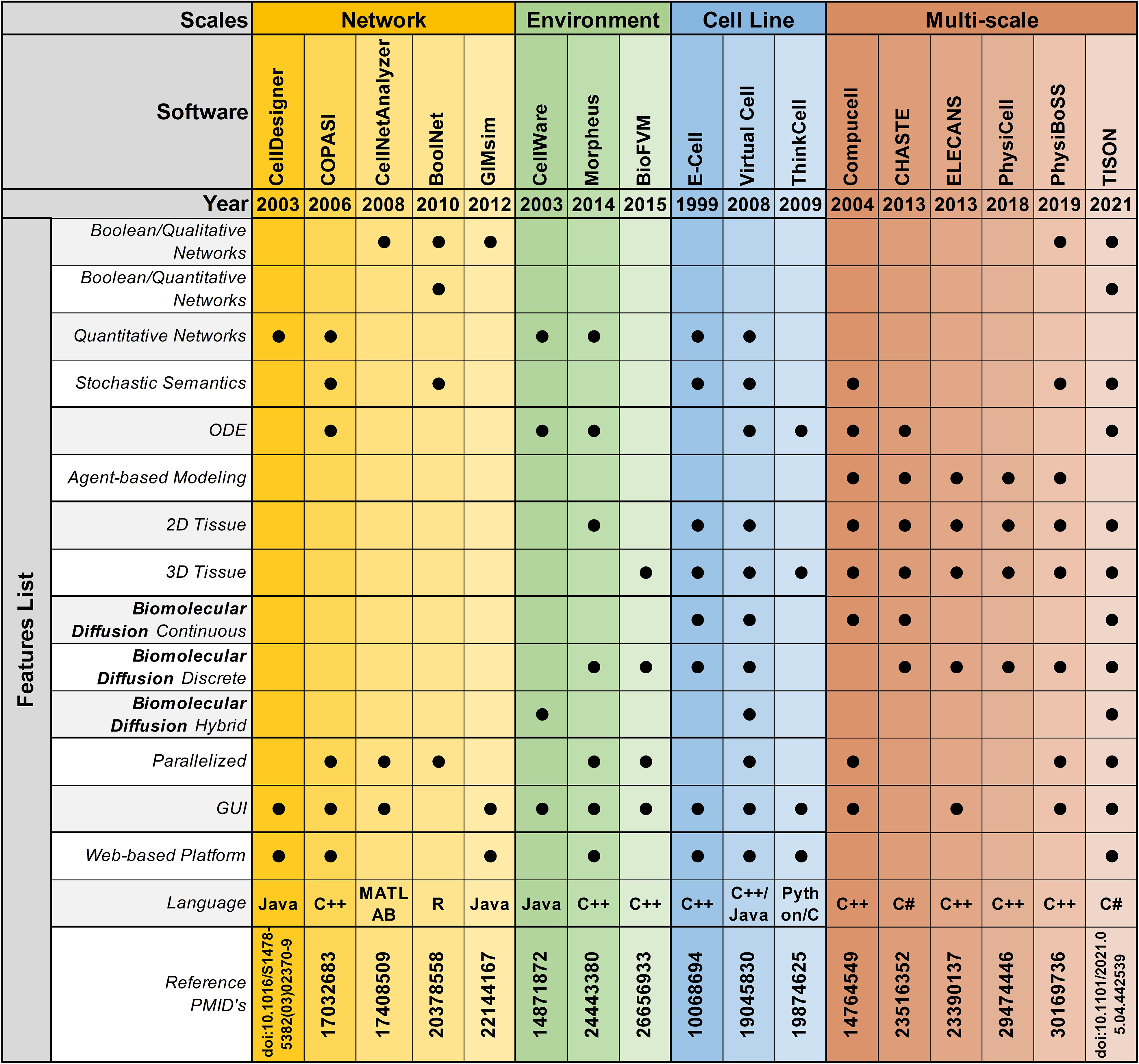
Feature-by-feature comparison of networks, environments, cell lines, and multi-scale modeling software in chronological order.

#### Biomolecular Modeling Platforms

Software platforms aimed at modeling biomolecular entities, abstract information from published literature as well as high-throughput technologies, and model them using a Boolean modeling approach or a differential equations modeling strategy.

#### Boolean Modeling Software

Boolean network modeling technique was first introduced by Kauffman (247, 248), in 1969. This approach has been widely adopted as a tool to model gene, transcript, protein, and metabolite regulatory networks. Numerous mathematical and computational cancer models have been developed using this representation (180,232–236). To facilitate the Boolean model development and analysis process, several platforms have been devised (249–251) (Supplementary Table 8). The applicability of these platforms can be further categorized into qualitative or quantitative Boolean networks modeling.

##### (i) Qualitative Boolean Modeling

Biomolecular qualitative Boolean models are a widely employed approach in cancer systems biology research to cater for cases where there is insufficient quantitative information, and/or lack of mechanistic understanding. Thus far, numerous platforms have been reported to help researchers develop qualitative Boolean network models. Amongst them, FluxAnalyzer (252), reported by Klamt *et al.* in 2003, was developed to undertake metabolic pathway construction, flux optimization, topological feature detection, flux analysis, etc. To expand the scope of the platform and include cell signaling, gene as well as protein regulatory networks, in 2007, Klamt *et al*. expanded FluxAnalyzer and developed CellNetAnalyzer (237). Tian *et al.* (253) employed CellNetAnalyzer to develop a p53 network model for evaluating DNA damage in cancer, while Hetmanski *et al.* (254) designed MAPK-driven feedback loop in Rho-A-driven cancer cell invasion. Although CellNetAnalyzer remains a widely used logical modeling software, however, its programmability and MATLAB dependency hinders its clinical employment towards the development of personalized cancer models. Towards addressing this challenge, in 2008, Albert *et al.* reported BooleanNet (250), an open-source, freely available Boolean modeling software for large-scale simulations of dynamic biological systems. Saadatpoort *et al.* (255) employed BooleanNet’s general asynchronous (GA) method to deduce therapeutic targets for granular lymphocyte leukemia. Similarly, in 2008, Kachalo *et al* reported NET-SYNTHESIS (238); a platform for undertaking network synthesis, inference, and simplification. Steinway *et al.* (256) employed both BooleanNet and NET-SYNTHESIS platforms to model epithelial-to-mesenchymal transition (EMT) in light of TFGβ cell signaling, in hepatocellular carcinoma patients towards elucidating potential therapeutic targets. In particular, BooleanNet was used to undertake model simulation and NET-SYNTHESIS for carrying out network interference and simplification. Another logical modeling platform, GIMsim (257), published by Naldi *et al.,* in 2009, also employed asynchronous state transition graphs to perform qualitative logical modeling which is especially useful for networks with large state space. This platform was employed by Flobak *et al.* (258) to map cell fate decisions in gastric adenocarcinoma cell-line towards evaluating drug synergies for treatment purposes, while Remy *et al.* (259) studied mutually exclusive and co-occurring genetic alterations in bladder cancer. GIMsim also implemented multi-valued logical functions, useful in simulating qualitative dynamical behavior in cancer research. However, the platform was unable to program automatic theoretical predictions, moreover, it only employed qualitative analysis approaches and could not be used to accurately map cell fates based on quantitative biomolecular expression data. Taken together, classical qualitative Boolean modeling approaches remain limited in developing predictive cancer models that could leverage quantitative biomolecular expression data generated from next-generation proteomics and related-sequencing projects.

##### (ii) Quantitative Boolean Modeling

Platforms aimed to integrate quantitative expression data from existing literature and databases towards carrying out network annotation and onwards analysis can be particularly useful in developing personalized cancer models. One such platform, the Markovian Boolean Stochastic Simulator (MaBoss), was established by Stoll *et al.* (260) in 2017, for stochastic and semi-quantitative Boolean network model development, mutations, and drug evaluation, sensitivity analysis based on experimental data, and eliciting model predictions. In 2019, Béal *et al.* (198) employed MaBoss to develop a logical model to evaluate breast cancer in light of individual patients’ genomic signature for personalized cancer medicine. This model was later expanded, in 2021, to investigate BRAF treatments in melanomas and colorectal cancer patients (199). Similarly, Kondratova *et al*. (261) used MaBoss to model an immune checkpoint network to evaluate the synergistic effects of combined checkpoint inhibitors in different types of cancers. In a similar attempt at developing quantitative Boolean networks, BoolNet (262) was reported in 2010 by Müssel and Kestler. BoolNet allows its users to reconstruct networks from time-series data, perform robustness and perturbation analysis and visualize the resultant cell fates attractor. BoolNet was employed by Steinway *et al*. (263) to construct a metabolic network towards evaluating gut microbiome in normal and disease conditions, whereas, Cohen *et al.* (264) studied tumor cell invasion and migration. BoolNet, however, lacks a graphical user interface, and results from the analysis cannot be visualized interactively, which hindered its employment. To address this issue, Shah *et al*. (265), in 2018, developed an Attractor Landscape Analysis Toolbox for Cell Fate Discovery and Reprogramming (ATLANTIS). ATLANTIS has an intuitive graphical user interface and interactive result visualization feature, for ease in utilization. The platform can be employed to perform deterministic as well as probabilistic analysis and was validated through literature-based case studies on the yeast cell cycle (266), breast cancer (267), and colorectal cancer (268).

#### Differential Equations Modeling Software

Although Boolean models have proven to be a powerful tool in modeling complex biomolecular signaling networks, however, these models are unable to describe continuous concentration, and cannot be used to quantify the time-dependent behavior of biological systems, necessitating the need to switch to quantitative differential equations (269). As a result, numerous stand-alone and web-based tools have been developed to build continuous network models to help describe the temporal evolution of biomolecules towards elucidating more accurate cell fate outcomes from quantitative expression data (Supplementary Table 8). Amongst initial attempts at developing such software, GEPASI (GEneral PAthway Simulator) was reported by Pedro Mendes *et al.* (239), in 1993. GEPASI is a stand-alone simulator that facilitates formulating mathematical models of biochemical reaction networks. GEPASI can also be used to perform parametric sensitivity analysis using an automatic pipeline that evaluates networks in light of exhaustive combinatorial input parameters. Ricci *et al.* (270) employed GEPASI to investigate the mechanism of action of anticancer drugs, while Marín-Hernández *et al.* (271) constructed kinetic models of glycolysis in cancer. In 2006, Hoops *et al*. reported a successor of GEPASI; COPASI (COmplex PAthway SImulator) (272), a user-friendly independent biochemical simulator that can handles larger networks for faster simulation results, through parallel computing. Orton *et al.* (273) employed COPASI to model cancerous mutations in EGFR/ERK pathway, while cellular senescence was evaluated by Pezze *et al.* (274) for targeted therapeutic interventions. Towards establishing a user-friendly software with an intuitive graphical user interface (GUI), another desktop application, CellDesigner was published by Funahashi *et al.* (275). CellDesigner application can be extended to include various simulation and analysis packages through integration with systems biology workbench (SBW) (276). In a case study using CellDesigner, Calzone *et al.* (277) developed a network of retinoblastoma protein (RB/RB1) and evaluated its influence in the cell cycle, while Grieco *et al.* (278) investigated the impact of Mitogen-Activated Protein Kinase (MAPK) network on cancer cell fate outcomes.

#### Environmental Modeling Software

To facilitate the development of environmental models that can help investigate inter-, intra-, and extracellular interactions between cellular network models and their dynamic environment, several software and platforms have been reported (240, 279). These platforms employ discrete, continuous, and hybrid approaches to develop models of cellular microenvironments towards setting up specific biological contexts such as normoxia, hypoxia, Warburg effect, etc (Supplementary Table 9). In 2014, Starruß *et al.* (240) published Morpheus, a platform for modeling complex tumor microenvironment. Morpheus leverages a cellular potts modeling approach to integrate and stimulate cell-based biomolecular systems for modeling intra- and extra cellular dynamics. In a case study using Morpheus, Felix *et al.* (280) evaluated pancreatic ductal adenocarcinoma’s adaptive and innate immune response levels, while Meyer *et al*. (281) investigated the dynamics of biliary fluid in the liver lobule. Although Morpheus is a widely employed modeling software, however, its diffusion solver is limited in its capacity to model large 3D domains. Towards modeling fast simulations for larger cellular systems, the Finite Volume Method for biological problems (BioFVM) software (279) was reported by Ghaffarizadeh *et al.*, in 2016. BioFVM is an efficient transport solver for single as well as multi-cell biological problems such as excretion, decomposition, diffusion, and consumption of substrates, etc (282). In a case study using BioFVM, Ozik *et al.* (283) evaluated tumor-immune interactions, while Wang *et al.* (284) elucidated the impact of tumor-parenchyma on the progression of liver metastasis. BioFVM, however, relies on its users to have programming-based knowledge to develop their models, which limits its translational potential. Towards minimizing programming requirement, SALSA (ScAffoLd SimulAtor) was reported by Cortesi *et al*. (218) in 2020. SALSA is general-purpose software that employs a minimum programming requirement, a significant advantage over its predecessors. The platform, in particular, has been useful in studying cellular diffusion in 3D cultures. This recent tool was validated in 2021, with a case study that evaluated and predicted therapeutic agents in 3D cell cultures (285).

#### Cell-level Modeling Software

Towards modeling cancer cell-specific behaviors such as cellular adhesion, membrane transport, loss of cell polarity, etc several software have been reported which can help develop *in silico* cancer cellular models (Supplementary Table 10). The foremost endeavor to develop software for cell-level modeling and simulation, led to the establishment of E-Cell (241), in 1999. E-Cell can be employed to model biochemical regulations and genetic processes using biomolecular regulatory networks in cells. In a case study using E-Cell, Edwards *et al*. (286) predicted the metabolic capabilities of *Escherichia coli*, and validate the results using existing literature, while Orton *et al*. (287) modeled the receptor-tyrosine-kinase-activated MAPK pathway. This software was expanded in 2001, with the development of Virtual Cell (V-Cell) (242,243,288), a web-based general-purpose modeling platform. V-Cell has an intuitive graphical and mathematical interface that allows ease in the design and simulation of whole cells, along with sub-cellular biomolecular networks and the external environment. Neves *et al*. (289) employed V-Cell to investigate the flow of spatial information in cAMP/PKA/B-Raf/MAPK1,2 networks, while Calebiro *et al*. (290) modeled cell signaling by internalized gprotein-coupled receptors. Similarly, to increase the efficiency in the design and modeling of synthetic regulatory networks in cells, in 2009, Chandran *et al*. reported TinkerCell (291) with computer-aided design (CAD) functionality which enabled faster simulation and associated analysis. The platform employed a modular approach for constructing networks and provides built-in features for ease in network construction, robustness analysis, and evaluating networks using existing databases. The evolutionary trend for TinkerCell’s platform adaptability and flexibility is highlighted in Figure 4, which shows a gradual shift from being a model specific platform to a domain modelling one. In a study employing TinkerCell, Renicke *et al*. (292) constructed a generic photosensitive degron (psd) model to investigate protein degradation and cellular function, while Chandran *et al*. (293) reported computer-aided biological circuits.

**Figure 4.**
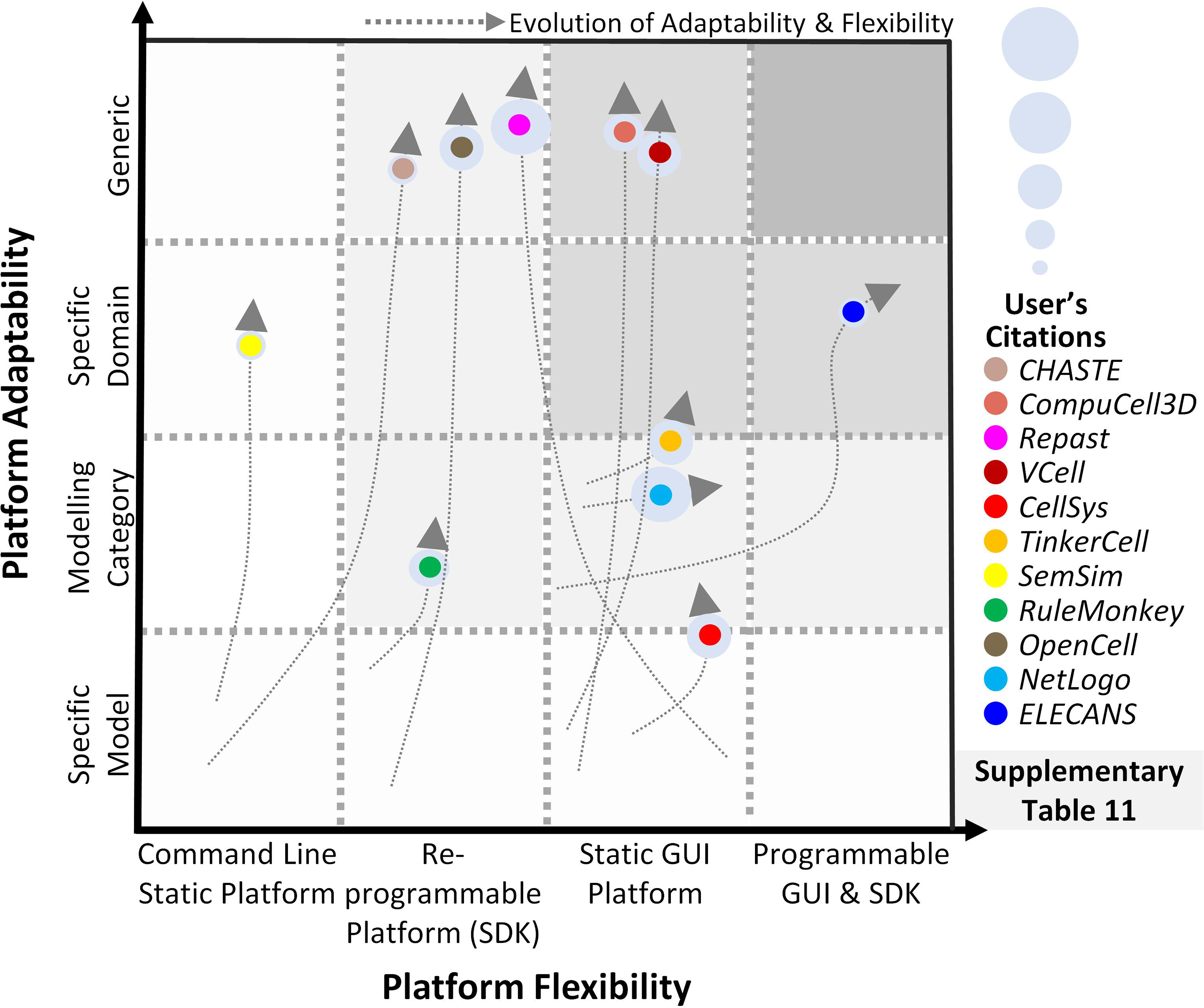
Evolution of scale-specific and multi-scale software. Evolution of multiscale modeling software for abstracting and simulating the spatiotemporal biomolecular complexity. Highlighting the need for a generic, data-driven, zero-code software requirement.

#### Multi-scale Modeling Software

Multi-scale cancer modeling approaches bring together scale-specific models towards undertaking an integrative analysis of heterogeneous experimental data by building coherent and biologically plausible models (294–297). Several multi-scale modeling platforms have been reported to help develop multiscale cancer models (244–246) (Supplementary Table 11). Amongst these, the REcursive Porous Agent Simulation Toolkit (Repast) (298, 299) published in 2003, provides a free and open-source tool for modeling and simulating agent-based models, with high-performance computing (HPC) capability. Repast toolkit was employed in 2007, by Folcik *et al.* (300) to develop an agent-based model which was then used to study interactions between cells and the immune system. Similarly, Mehdizadeh *et al*. (301) used Repast to model angiogenesis in porous biomaterial scaffolds. Although Repast can be used for simulating several types of evolutionary trends between agents, there is no established guideline for selecting a mechanism to model such trends, hindering its use by naïve users. Moreover, Repast does not have a GUI or a software development kit (SDK) interface for implementing subcellular mechanisms e.g., gene, protein, and metabolic networks. In contrast, CompuCell (295), published in 2004 by Izaguirre *et al.*, provides an elaborative GUI to model cell-scale or tissue-scale simulations by integrating biomolecular networks, intra- and extracellular environment, and cell to environment interactions. Mahoney *et al.* (302) employed CompuCell to develop an angiogenesis-based model in cancer for investigating novel cancer therapies. This model was later optimized in 2010 (303) for evaluating potential new approaches for therapeutic intervention for angiogenesis. Although CompuCell provides an intuitive framework modeling paradigm, however, the platform core is not conducive to multi-scale cancer modeling. The focus of the software is primarily multi-agent simulations rather than multi-scale cancer modeling. This poses a significant challenge in the utilization of the software. In 2013, CHASTE (296) (Cancer Heart and Soft Tissue Environment) was launched, which provides a computational simulation pipeline for mathematical modeling of complex multi-scale models. Users can employ CHASTE for a wide range of problems involving on and off-lattice modeling workflows. CHASTE has also previously been employed to model colorectal cancer crypts. Nonetheless, CHASTE does not have a GUI and can only be executed by command line text commands. Furthermore, it requires recompilation on part of the modeler to use the code updates performed by the group. To further improve the multi-scale modeling approach for investigating cancer, in 2013, Chaudhary *et al*. (297) published ELECANS (Electronic Cancer System). ELECANS had an intuitive but rigid GUI along with a programmable SDK besides the lack of a high-performance simulation engine. ELECANS was employed to model the mitochondrial processes in cancer towards elucidating the hidden mechanisms involved in the Warburg effect (304). Although ELECANS provided a feature-rich environment for constructing multi-scale models, however, the platform lacked a biomolecular network integration pipeline, and it also placed a heavy programming requirement on its users. In contrast, in 2018, PhysiCell (305) was reported for 2D and 3D multi-cell off-lattice agent-based simulations. PhysiCell is coupled with BioFVM (279)’s finite volume method to model multi-scale cancer systems (283). In a salient example, Wang *et al.* (284) employed PhysiCell to model liver metastatic progression. In 2019, PhysiCell’s agent-based modeling features and MaBoss’s Boolean cell signaling network feature were coupled together to develop an integrated platform, PhysiBoss (306). As a result, PhysiBoss provided an agent-based modeling environment to study physical dimension and cell signaling networks in a cancer model. In 2020, Colin *et al*. (307) employed PhysiBoss’s source code to model diffusion in oocytes during prophase 1 and meiosis 1, while Getz *et al*. (308) proposed a framework using PhysiBoss to develop a multi-scale model of SARS-CoV-2 dynamics in lung tissue. Recently, in 2021, Gondal *et al*. (309) reported Theatre for *in silico* Systems Oncology (TISON), a web-based multi-scale “zero-code” modeling and simulation platform for *in silico* oncology. TISON aims to develop single or multi-scale models for developing personalized cancer therapeutics. To exemplify the use case for TISON, Gondal *et al*. employed TISON to model colorectal tumorigenesis in *Drosophila melanogaster’*s midgut towards evaluating efficacious combinatorial therapies for individual colorectal cancer patients (236).

Summarily, multi-scale modeling software have enabled the development of biologically plausible cancer models to varying degrees. These platforms, however, fall short of providing a generic and high-throughput environment that could be conveniently translated in clinical settings.

### Pipelining Panomics Data towards *in silico* Clinical Systems Oncology

*In silico* multi-scale cancer models, annotated with patient-specific biomolecular and clinical data, can help decode complex mechanisms underpinning tumorigenesis and assist in clinical decision-making (310, 311). Clinically-driven *in silico* multi-scale cancer models simulate *in vivo* tumor growth and response to therapies across biocomplexity scales, within a clinical environment, towards evaluating efficacious treatment combinations. To facilitate the development of multi-scale cancer models, several large-scale program projects have been launched (Supplementary Table 12) such as Advancing Clinico-Genomic Trials on Cancer (ACGT) (312), Clinically Oriented Translational Cancer Multilevel Modeling (ContraCancrum) (313), Personalized Medicine (p-medicine) (314), Transatlantic Tumor Model Repositories (TUMOR) (315), and Computational Horizons In Cancer (CHIC) (316), amongst others (Figure 5). Here, we review and evaluate five salient projects for multi-scale cancer modeling towards their clinical deployment.

**Figure 5.**
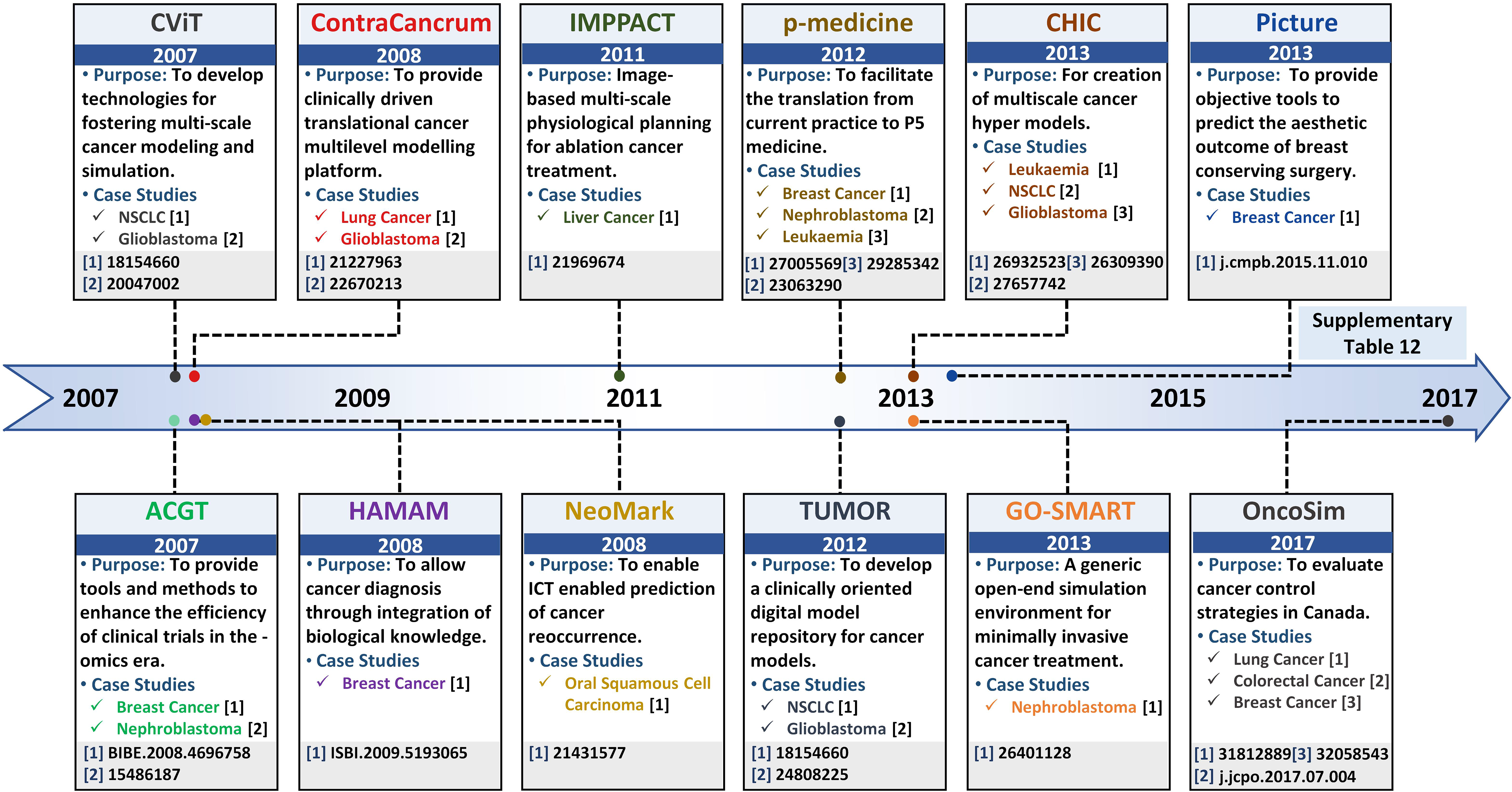
Salient projects pipelining multi-scale panomics data into clinical settings – a timeline. Timeline highlighting salient project platforms for developing realistic and clinically-driven multi-scale cancer models, along with their associated leading case studies.

#### ACGT Project

The ACGT project (312), launched in 2007, proposed to develop Clinico-Genomic infrastructure for organizing clinical and genomic data towards investigating personalized therapeutics regimens for an individual cancer patient (317). The ACGT platform provides an open-source and open-access infrastructure designed to support the development of “oncosimulators” to help clinicians accurately compare results from different clinical trials and enhance their efficiency towards optimizing cancer treatment (318, 319). The ACGT framework employs molecular and clinical data generated from different sources including whole genome sequencing, histopathological, imaging, molecular, and clinical data, etc. to develop simulators for mimicking clinical trials. Personalized panomics data employed to develop oncosimulators in ACGT is extracted from real patients which enables the oncosimulators to be clinically relevant for predictive purposes. Additionally, ACGT provides data retrieval, storage, integrative, anonymization, and analysis as well as results presentation capabilities. Using the platform, in 2008, Kolokotroni *et al*. (320) developed a personalized, spatiotemporal oncosimulator model of breast cancer to mimic a clinical trial based on protocols outlined in the Trail of Principle (TOP), towards evaluating the model response to chemotherapeutic treatment in neoadjuvant settings (321). Similarly, in 2009, Graf *et al*. (322, 323) modeled nephroblastoma oncosimulator, a childhood cancer of the kidney, based on a clinical trial run by the International Society of Paediatric Oncology (SIOP) for simulating tumor response to therapeutic regimens in the clinical trials. The results generated from the TOP and SIOP trials enabled the ACGT oncosimulators to adapt in light of real clinical conditions and the software to be validated against multi-scale patient data. The focus of ACGT oncosimulators, however, is limited to existing clinical trials for predicting efficacious treatment combinations.

#### The ContraCancrum Project

Towards establishing a platform for the development of composite multi-scale models for simulating malignant tumor models, in 2008, ContraCancrum project (313) was initiated. ContraCancrum aimed to develop a multi-scale computational framework for translating personalized cancer models into clinical settings towards simulating malignant tumor development and response to therapeutic regimens. For that, the Individualized MediciNe Simulation Environment (IMENSE) platform (324, 325) was established, to undertake the oncosimulator development process. The platform was employed for molecular and clinical data storage, retrieval, integration, and analysis. Oncosimulators, developed under the ContraCancrum project, employ patient data across biologically and clinically relevant scales including molecular, environmental, cells, and tissues level. These oncosimulators can be used to optimize personalized cancer therapeutics for assisting clinician decision-making process. To date, several oncosimulators have been developed, under the ContraCancrum project (31,313,323,325–332). In particular, the initial validation of the ContraCancrum workflow was performed using two case studies; glioblastoma multiforme (GBM) (313,326,332) and non-small cell lung cancer (NSCLC) (313,330,331). In 2010, Folarin and Stamatakos (333) developed a glioblastoma oncosimulator using personalized molecular patient data to evaluate treatment response under the effect of a chemotherapeutic drug (temozolomide) (324). Similarly, Roniotis *et al*. (332) developed a multi-scale finite elements workflow to model glioblastoma growth, while Giatili *et al*. (326) outlined explicit boundary condition treatment in glioblastoma using an *in silico* tumor growth model. The results from the case studies were validated by comparing *in silico* prediction with pre- and post-operative imaging and clinical data (313, 328). Moreover, The ContraCancrum project hosts more than 100 lung patients’ tumor and blood samples (313). This data is employed to develop clinically validated *in silico* multilevel cancer models for NSCLC using patient-specific data (330). In 2010, using a biochemical oncosimulator, Wan *et al*. (330) investigated the binding affinities for AEE788 and Gefitinib tyrosine kinase inhibitor against mutated epidermal growth factor receptor (EGFR) for NSCLC treatment. Similarly, Wang *et al*. (331) developed a 2D agent-based NSCLC model to investigate proliferation markers and evaluated ERK as a suitable target for targeted therapy. Although ContraCancrum’s project has created numerous avenues for multilevel cancer model development, however, the platform is limited to only specific types of cancer for which data is internally available on the platform.

#### p-medicine Project

Towards improving clinical deployment capabilities of oncosimulators, another pilot project: the personalized medicine (p-medicine) project was launched in 2011 (314). The main aim of p-medicine was to create biomedical tools facilitating translation of current medicine to P5 medicine (predictive, personalized, preventive, participatory, and psycho-cognitive) (334). For that, the p-medicine portal provides a web-based environment that hosts specifically-purpose tools for personalized panomics data integration, management, and model development. The portal has an intuitive graphical user interface (GUI) with an integrated workbench application for integrating information from clinical practices, histopathological imaging, treatment and omics data, etc. Computational models developed under p-medicine workflow, are quantitatively adapted to clinical settings since they are derived using real multi-scale data. Several multi-scale cancer simulation models (oncosimulators) (314, 334) have been devised, using the p-medicine workflow. Amongst these, in 2012, Georgiadi *et al*. (335) developed a four-dimensional nephroblastoma treatment model and evaluated its employment in clinical decisions making. Towards evaluating personalized therapeutic combinations, in 2014, Blazewicz *et al*. (336) developed a p-medicine parallelized oncosimulator which evaluated nephroblastoma tumor response to therapy. The parallelization enhanced model usability and accuracy for eventual translation into clinical settings for supporting clinical decisions. In 2016, Argyri *et al*. (337) developed a breast cancer oncosimulator to evaluate vascular tumor growth in light of single-agent bevacizumab therapy (anti-angiogenic treatment), while in 2014, Stamatakos *et al*. (338) evaluated breast cancer treatment under an anthracycline drug for chemotherapy (epirubicin). In 2012, Ouzounoglou *et al.* (339) developed a personalized acute lymphoblastic leukaemia (ALL) oncosimulator for evaluating the efficacy of prednisolone (a steroid medication). This study was further augmented, in 2015, by Kolokotroni *et al*. (340) to investigate the potential cytotoxic side effects of prednisolone. Later in 2017, Ouzounoglou *et al.* (341) expanded their ALL oncosimulator model to a hybrid oncosimulator for predicting pre-phase treatments for ALL patients. The validation of these models was undertaken using clinical trial data in pre and post-treatment (342). Although, p-medicine presents a state-of-the-art *in silico* multi-scale cancer modeling environment, however, the project is limited in its application for determining individual biomarkers for potential novel therapy identification. Moreover, the project is limited to its niche set of tools and models that cannot be integrated with other similar model repositories and platforms.

#### TUMOR Project

In 2012, a transatlantic USA-EU partnership was initiated with the launch of Transatlantic Tumor Model Repositories (TUMOR) (315). TUMOR provides an integrated, interoperable transatlantic research environment for developing a clinically-driven cancer model repository. The repository aimed to integrate computational cancer models developed by different research groups. TUMOR’s transatlantic aim was to couple models from ACGT and ContraCancrum projects with those at the Center for the Development of a Virtual Tumor (CViT) (343) as well as other relevant centers. TUMOR is predicted to serve as an international clinical translation, interoperable and validation platform for *in silico* oncology hosting multi-scale cancer models from different cancer model repositories and platforms such as E-Cell (344), CellML (345), FieldML (346), and BioModels (39, 347). To support model data interoperability between platforms, TumorML (Tumor model repositories Markup Language) (347) was developed towards facilitating inter-data operability in the TUMOR project. The TUMOR environment also offers a wide range of additional services supporting predictive oncology and individualized optimization of cancer treatment. For example, the platform allows remote access of predictive cancer models in hospitals and to clinicians for the development of quantitative cancer research and personalized cancer therapy model. TUMOR environment also incorporates deterministic as well as stochastic models through COPASI simulator (246). An automatic validation pipeline is also embedded for the execution and deployment of these models in clinical settings (315). TUMOR’s ability to couple and integrate models from different scales, approaches, and repositories towards increasing model accuracy in predictive oncology was exemplified with Wang *et al*.’s (348) NSCLC model. In 2007, Wang *et al*. developed a 2D multi-scale NSCLC model for evaluating tumor expansion dynamics in NSCLC patients, in CViT. This model was exported in TumorML and made available in the TUMOR repository to help evaluate growth factor influence in aggressive cancer (349). Similarly, in 2014 Sakkalis *et al*. (350) employed the TUMOR platform to interlink and couple three independent glioblastoma-specific cancer models (EGFR signaling (351), cancer metabolism (352), Oncosimulator (350, 353)), reported by different research groups. The resultant model was used to investigate the impact of radiation and temozolomide (chemotherapy) on glioblastoma multiforme to evaluate treatment effectiveness.

#### CHIC Project

Another transatlantic project Computational Horizons In Cancer (CHIC) project (316) launched in 2013, aimed to provide an oncosimulator modeling platform for *in silico* oncology. CHIC was initiated to develop and implement predictive oncology and individualized multi-scale cancer modeling tools towards assisting quantitative cancer research and personalized cancer therapy. The workflow and tools established under the ambient of the CHIC project allowed the development of robust, interoperable, and collaborative *in silico* models in cancer and normal conditions (316). The CHIC project also proposed a pipeline to translate the model towards supporting clinicians to make optimal personalized treatment plans for individual patients. To this end, several models are created using CHIC such as non-small cell lung cancer (354), glioblastoma (355), leukemia model (356), etc. These models furnish a quantitative understanding of tumorigenesis towards providing avenues for promoting individual cancer patient treatment combinations. One such model reported by Kolokotroni *et al*. (354) evaluated the efficacy of cisplatin-based therapy for NSCLC patients using *in silico* multi-scale cancer model. While, in 2015, Antonopoulos and Stamatakos (355) modeled the infiltration of glioblastoma cells in normal brain regions using a novel treatment. In 2017, Stamatakos and Giatili (357) extended glioblastoma oncosimulator for modeling tumor growth using reaction-diffusion numerical handling based on multi-scale panomics data. They further proposed a clinical pipeline to translate the model into clinical settings towards supporting clinicians to make optimal personalized treatment plans for individual patients. Ouzounoglou *et al*. (356) developed an *in silico* multi-scale leukemia oncosimulator model towards modeling deregulations in the G1/S pathway to investigate the altered function of retinoblastoma in ALL patients. However, a clinical translation of these models is currently in the works (358).

## Future Directions and Conclusion

In this work, we have evaluated the use of data-driven multi-scale cancer models in deciphering complex biomolecular underpinnings in cancer research towards developing personalized therapeutics interventions for clinical decision-making. Specifically, we have discussed the chronological evolution of online cancer data repositories that host high-resolution datasets from multiple spatiotemporal scales. Next, we evaluate how this data drives single- and multi-scale systems biology models towards decoding complex cancer regulation in patients. We then track the development of various modeling software and their applications in enhancing the translational role of cancer systems biology models in clinics. We conclude that the contemporary multi-scale modeling software line-up remains limited in their clinical employment due to the lack of a generic, zero-code, panomic-based framework for translating research models into clinical settings. Such a framework would help annotate *in silico* cancer models developed using single and multi-scale databases (69,359–365). The framework should also provide an environment for developing the extra-cellular matrix of a cancer cell which can then be integrated into cellular models. Existing environment (42, 43) and cell line databases (45–47) need to be integrated to design environmental models along with biologically plausible cell line structures. These cell lines could then be assembled into three-dimensional geometries to create multi-scale *in silico* organoids. The pipeline should also furnish capabilities such as a convenient import workflow for clinical data integration through histopathological image data repositories (48–50) for designing biologically accurate organoid structures based on each cancer patient’s underlying cellular morphology. Once the personalized multi-scale model has been constructed, the pipeline would allow investigation into the temporal evolution of the multi-scale organoid under personalized inputs and user-designed biomolecular entities. Data generated from the multi-scale model simulation can be analyzed to elicit biomolecular cues for each cancer patient as well as determine its role.

Taken together, a translational *in silico* systems oncology pipeline is the need of the hour and will help develop and deliver personalized treatments of cancer as well as substantively inform clinical decision-making processes.

## Conflict of Interest

The authors declare that they have no conflict of interest.

## Funding

This work was supported by the National ICT-R&D Fund (SRG-209), RF-NCBC-015, NGIRI-2020-4771, HEC (21-30SRGP/R&D/HEC/ 2014, 20-2269/NRPU/R&D/ HEC/12/ 4792 and 20-3629/NRPU/R&D/HEC/14/ 585), TWAS (RG 14-319 RG/ITC/AS_C) and LUMS (STG-BIO-1008, FIF-BIO-2052, FIF-BIO-0255, SRP-185-BIO, SRP-058-BIO and FIF-477-1819-BIO) grants.

## Data Availability Statement

The supplementary tables for the manuscript titled “*Navigating Multi-scale Cancer Systems Biology towards Model-driven Personalized Therapeutics*” is available online.

